# Extracellular HSPB1 protects intestinal epithelial cells against ethanol-activated ferroptosis

**DOI:** 10.64898/2026.10.14.759219

**Authors:** McKenzie Burge, Anne-Marie C Overstreet, B Ben Koff, Steven H. Overend, Kylee A. Hunter, Jeannette S Messer

**Affiliations:** Department of Microbial Sciences in Health, Cleveland Clinic Research, Cleveland Clinic, Cleveland, OH, USA; Department of Molecular Medicine, Cleveland Clinic Lerner College of Medicine of Case Western Reserve University, Cleveland, OH, USA; Northern Ohio Alcohol Center, Lerner Research Institute, Cleveland Clinic, Cleveland, Ohio, USA; Center for Microbiome and Human Health, Lerner Research Institute, Cleveland Clinic, Cleveland, Ohio, USA

**Keywords:** Alcohol, ethanol toxicity, oxidative damage, iron, damage-associated molecular pattern molecule, gastrointestinal tract, gut-liver axis

## Abstract

Alcohol consumption damages the gastrointestinal tract, the initial site of contact for ingested ethanol. Intestinal epithelial cells (IEC) lining the small intestine are at particular risk of ethanol toxicity since they are largely responsible for ethanol absorption and metabolize ethanol into toxic metabolites. These toxic metabolites generate oxidative stress that damages cellular components and leads to programmed cell death. The small heat shock protein HSPB1 regulates oxidative stress and provides cytoprotection in cells exposed to ethanol, but its specific role in intestinal ethanol toxicity is still unclear. Therefore, the goal of this study was to understand the role of HSPB1 in IEC stress response pathways during ethanol exposure. Here we utilized *in vitro* models to demonstrate that IEC rapidly upregulate and release HSPB1 in response to ethanol. HSPB1 expression in IEC mitigated oxidative stress, oxidative damage, and iron uptake, the hallmarks of ferroptosis, during ethanol stress. Interestingly, recombinant HSPB1 rescued cells, suggesting that extracellular HSPB1 is responsible for its protective effect. Our findings establish extracellular HSPB1 as a previously unrecognized mediator of IEC cytoprotection during acute ethanol exposure. However, HSPB1 functions as a “double-edged sword” along the gut-liver axis. While transient extracellular HSPB1 protects the gut epithelium from acute oxidative damage, sustained levels driven by chronic alcohol consumption lead to HSPB1 release from hepatocytes that activates macrophages and contributes to driving self-damaging inflammation in conditions like alcohol-associated hepatitis. Therefore, targeting extracellular HSPB1 levels along the gut-liver axis offers a promising diagnostic and therapeutic avenue to mitigate severe alcohol-induced organ injury.

## INTRODUCTION

Chronic alcohol consumption causes damage to organs throughout the body.(1) However, the gastrointestinal (GI) tract is the point of first contact between alcohol and human tissue after ingestion. Within the GI tract, alcohol directly and indirectly damages the epithelial cells lining the oral cavity, stomach, and small and large intestines.(2) The small intestine is particularly vulnerable to injury since it absorbs a large portion of the ingested alcohol. Alcohol directly damages intestinal epithelial cell (IEC) membranes and its metabolism generates oxidative stress within these cells.(3) High levels of oxidative stress compromise IEC viability, alter cell metabolism, and lead to failure of the junctional complexes that tightly connect IEC and separate the contents of the gut lumen from the rest of the body. Additionally, alcohol damages microbes within the GI tract, leading to release of pro-inflammatory microbe-associated molecular patterns (MAMPs) and changes to the composition and function of the gut microbial community.(4, 5) These alcohol-induced changes within the GI tract also endanger the rest of the body since intestinal damage can compromise the GI barrier and allow MAMPs, such as lipopolysaccharide, to cross into portal circulation.(6) This means that protecting IEC from the damaging effects of alcohol limits damage to the GI tract and organs throughout the body. However, how IEC protect themselves from alcohol-induced damage is still not entirely clear.

Ethanol is the intoxicating component of alcoholic beverages. While the majority of ingested ethanol is metabolized in the liver, small intestinal epithelial cells also possess alcohol dehydrogenase, catalase, and the cytochrome P450 2E1 (CYP2E1) microsomal ethanol oxidizing system (MEOS).(7) These enzymes metabolize ethanol into the reactive metabolites acetaldehyde and NADP^+^, which cause oxidative stress that damages lipids, proteins, and DNA within cells. To counteract oxidative stress, IEC exposed to ethanol rapidly activate reactive oxygen species (ROS) mitigation systems that further metabolize acetaldehyde and NADP+ into non-harmful species, increase the concentration of antioxidants in the cell, and support repair of damaged cellular components. If these mitigation mechanisms are effective, IEC can clear the ROS and return to normal function. However, if they fail, IEC activate programmed cell death (PCD) mechanisms. The intestinal epithelium can compensate for and repair limited, localized IEC death, but extensive death can lead to mucosal damage, enteropathy, and barrier failure that allows MAMPs to escape the confines of the gastrointestinal tract lumen and enter portal circulation.(8) In the liver, MAMPs contribute to ethanol-activated inflammation which can become self-damaging if it is sustained long term.

HSPB1 (Heat Shock Protein Family B (Small) Member 1; also known as Hsp25/27) is a small heat shock protein that is rapidly induced in response to oxidative stress in many different cell types, including IEC. (9-11) During oxidative stress, HSPB1 acts as a molecular chaperone to refold damaged proteins, protects components of the cell cytoskeleton from damage, supports autophagy to clear protein aggregates and damaged organelles, and limits programmed cell death through direct interaction with proteins involved in apoptosis and ferroptosis.(12, 13) HSPB1 is reported to protect IEC from oxidative stress-activated death triggered by monochloramine and HSPB1 is induced in and released from hepatocytes exposed to ethanol.(14-16)

Evidence that HSPB1 protects IEC against oxidative stress and is activated in hepatocytes by ethanol led us to hypothesize that HSPB1 protects IEC against oxidative damage and death caused by exposure to ethanol. We tested this hypothesis in a series of experiments involving murine small intestine challenged with ethanol *ex vivo* and HSPB1 deficient human IEC exposed to ethanol *in vitro*. We found that HSPB1 is rapidly induced in response to ethanol and that it limits oxidative damage and iron accumulation in IEC, the hallmarks of ferroptosis. Despite the apparent failure of ROS-mitigation mechanisms, Nrf2 transcriptional activity was elevated in HSPB1-deficient IEC. Conversely, NF-kB activity, which also mitigates ROS-related damage, was decreased in the absence of HSPB1. HSPB1 was released from IEC challenged by ethanol and addition of extracellular HSPB1 rescued IEC from ethanol-activated death. Together, these findings establish a key role for extracellular HSPB1 in protection of the intestinal epithelium against damage caused by exposure to ethanol and suggest that it limits ethanol-triggered epithelial cell death by protecting cells against ferroptosis.

## RESULTS

### HSPB1 is rapidly increased in intestinal epithelial cells experiencing ethanol stress

HSPB1 is a redox-responsive molecular chaperone that protects cells against oxidative damage.(17) HSPB1 is normally maintained at low levels in unchallenged cells, but can be rapidly and robustly increased in response to oxidative stress, cytokine signals from nearby stressed cells, or MAMPs released from bacteria.(18) In IEC, HSPB1 preserves cellular viability in the presence of the strong oxidant monochloramine.(15) This led us to hypothesize that HSPB1 protects IEC from oxidative damage during ethanol exposure.

We first measured HSPB1 in murine small intestine exposed to ethanol *ex vivo*. This approach allowed us to compare levels of HSPB1 in unchallenged and challenged segments of small intestine from the same mouse. Exposure to a relatively low level of ethanol (1.5%) led to increased HSPB1 protein concentrations as early as 30 minutes after exposure (**Fig. 1A**). Since IEC lining the intestine are exposed to the highest levels of ethanol, we focused on HSPB1 in these cells. Similar to what we saw in the intestine, isolated IEC rapidly upregulated intracellular HSPB1 concentrations in response to ethanol and levels were sustained over several days with repeated ethanol exposure (**Fig. 1B, C, D**). Thus, IEC activate HSPB1 when they encounter ethanol.

**Figure 1.**
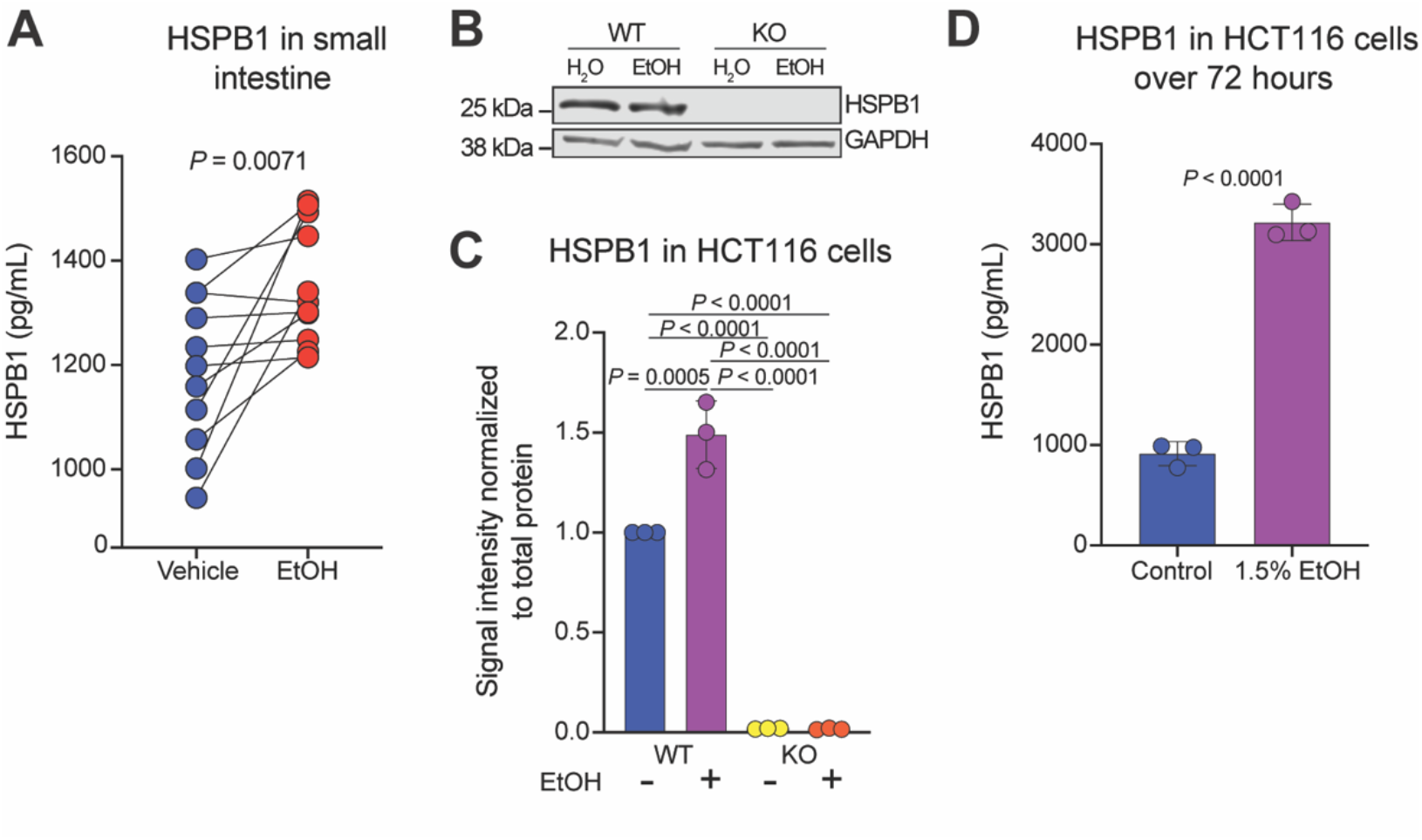
HSPB1 is rapidly increased in intestinal epithelial cells experiencing ethanol stress. **A.** HSPB1 concentration by ELISA in small intestine treated *ex vivo* with media or 1.5% ethanol for 30 min. Lines denote untreated (blue) and treated (red) sections from a single mouse (n=12; 6 male and 6 female). **B**. Immunoblot for HSPB1 and GAPDH in WT and KO HCT116 cells exposed to media or 1.5% ethanol for 1 hour. **C**. Quantification of band densitometry from B across three independent experiments. **D**. HSPB1 concentration by ELISA in WT HCT116 cells treated with media or 1.5% ethanol every 24 hr. for a total of 72 hr. Data plotted as mean + SD. (A) Paired *t* test. (D) Unpaired *t* test. (C) One-way ANOVA with a Tukey multiple comparisons post hoc test.

### HSPB1 protects IEC from ethanol-activated death

Ethanol can directly damage cells by acting on membrane lipids to compromise cell membrane integrity or it can indirectly damage cells through metabolism to reactive metabolites that damage components necessary for cellular function.(19) Each of these effects have the potential to compromise IEC viability and lead to cell death. For these reasons, we investigated the impact of HSPB1 on IEC survival during ethanol stress. Propidium iodide (PI) is a cell-membrane impermeant DNA dye. Cells with severely compromised cell membrane integrity allow PI to enter and bind to DNA, leading to increased fluorescence. In the absence of stress, PI fluorescence was similar between wild type (WT) and HSPB1 deficient (KO) IEC (**Fig. 2A, B**). However, in the presence of ethanol, KO cells had higher levels of PI uptake, indicating that these cells were no longer viable.

**Figure 2.**
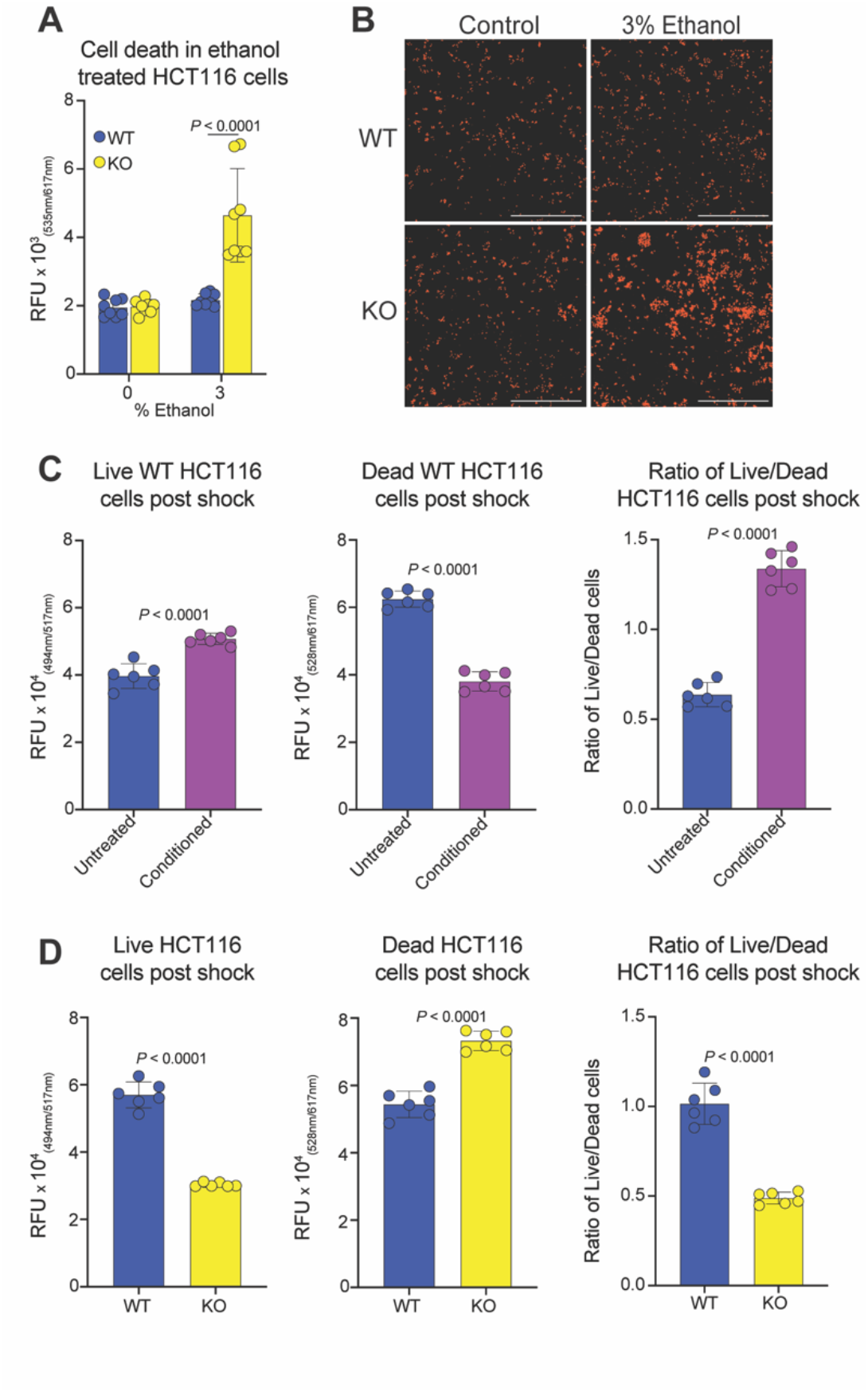
HSPB1 protects IEC from ethanol-activated death. **A.** Cell death in WT or KO HCT116 cells treated with media or 3% ethanol for 2 hours measured by PI. **B**. Microscope images of PI (orange) in samples from A. Scale bar = 1000 µm, 40X magnification. **C**. Live/Dead staining of WT HCT116 cells treated with 1.5% ethanol for 24 hr. then 3% ethanol for 1 hr. **C**. Live/Dead staining of WT or KO HCT116 cells treated with 1.5% ethanol for 24 hr. then 3% ethanol for 1 hr. Data plotted as mean + SD. (A,C,D) Unpaired *t* test.

Our finding that HSPB1 is increased by ethanol administration and remains high with repeated exposure led us to consider that HSPB1 could act as an ethanol conditioning factor. Conditioning is an effect of exposure to low levels of toxins that allows cells to survive higher levels of exposure than they would survive without conditioning.(20, 21) We first determined whether WT cells could be conditioned to increase survival during ethanol exposure. Live cells were identified with calcein-AM fluorescence that indicates intracellular esterase activity. Dead cells were identified by fluorescence with the cell impermeant dye ethidium homodimer-1 that indicates loss of cell membrane integrity. Conditioning with low level ethanol exposure protected WT cells from death caused by higher level ethanol exposure (**Fig. 2C**). In contrast, conditioning was not as effective in protecting KO cells from ethanol-activated death (**Fig. 2D**). Together, these findings indicate that activation of HSPB1 expression protects cells from ethanol-induced death.

### HSPB1 limits oxidative damage and iron accumulation during ethanol stress

Ethanol can trigger many different programmed cell death mechanisms, including apoptosis, necroptosis, pyroptosis, and ferroptosis.(22) Which cell death program ethanol activates is largely determined by the cell type, concentration and duration of ethanol exposure, and homeostatic conditions when a cell encounters ethanol. Previous studies have identified intracellular roles for HSPB1 in prevention of ferroptosis, a type of iron-dependent and non-apoptotic cell death that is activated in ethanol stress.(23-28) This led us to hypothesize that HSPB1 limits ferroptotic death in ethanol challenged IEC. We first measured reactive oxygen species (ROS) in WT and KO cells challenged by ethanol. The concentration of ROS increased in both cell types after ethanol challenge, but levels in ethanol stressed KO cells were nearly double those of WT cells (**Fig. 3A**). We next measured protein carbonylation, an indicator of oxidative damage, in WT and KO cells with and without ethanol stress. The level of carbonylated protein was not significantly increased in WT cells after ethanol exposure (**Fig. 3B**). However, the levels of carbonylated protein in KO cells were higher than WT at baseline and increased after ethanol stress. Protein carbonylation also serves as an indirect indication of lipid peroxidation. Oxidation of polyunsaturated fatty acids in cellular membranes generates lipid-derived electrophiles that react non-enzymatically with amino acid residues, this irreversible post-translational modification is known as protein carbonylation.(29) Finally, intracellular iron levels increased in WT cells with ethanol stress, but were more than double WT levels in the KO cells (**Fig. 3C**). The combination of oxidative stress, oxidative damage to proteins, and accumulation of intracellular iron shows that cells are dying of ferroptosis in our model. Our findings also show that HSPB1 protects IEC from this form of programmed cell death during ethanol stress.

**Figure 3.**
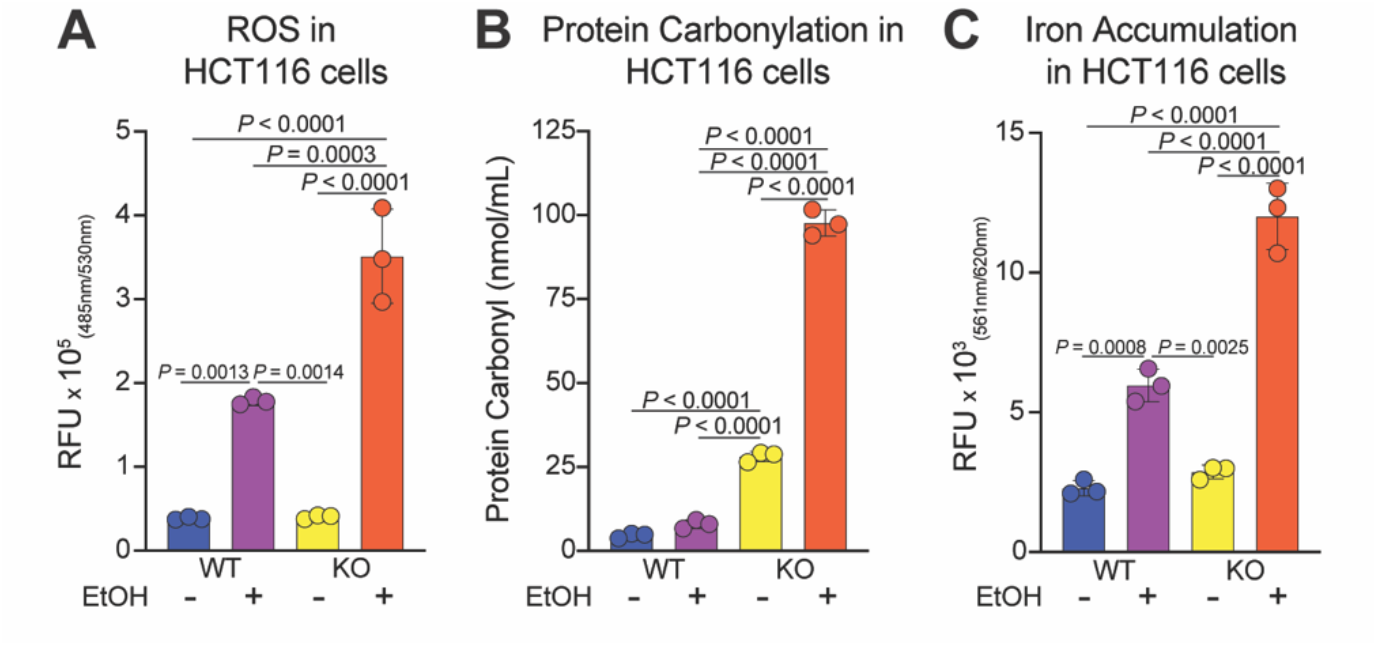
HSPB1 limits oxidative damage and iron accumulation during ethanol stress. **A.** Reactive oxygen species measured in WT or KO HCT116 cells treated with media or 1.5% ethanol for 1 hour. **B**. Protein carbonylation measured in WT or KO HCT116 cells treated with media or 1.5% ethanol for 1 hour. **C**. Iron accumulation measured in WT or KO HCT116 cells treated with media or 1.5% ethanol for 1 hour. Data plotted as mean + SD. (A,B,C) One-way ANOVA with a Tukey multiple comparisons post hoc test.

### HSPB1 is not required for activation of Nrf2 transcriptional activity, but is necessary for NF-kB activation during ethanol stress

Cells encountering ethanol rapidly activate oxidative stress response mechanisms to preserve cell functions and survive oxidative damage. The transcription factor Nrf2 (nuclear factor erythroid 2-related factor 2) is considered to be a master regulator of cellular defense against oxidative damage.(30) In unstressed cells, Nrf2 is trapped in the cytoplasm through association with Keap1 (Kelch-like ECH-associated protein 1). The association with Keap1 leads to constitutive proteasomal degradation of Nrf2, which maintains this vital protein at low levels, but in a state of readiness for a rapid response to oxidative stress. Generation of ROS leads to modification of cysteine residues in Keap1 which releases Keap1 from Nrf2 and allows Nrf2 to translocate to the nucleus where it activates a transcriptional program centered on production of antioxidants and cellular repair. A previous study identified HSPB1 as a critical factor in Nrf2 release from Keap1 during oxidative stress activated by hydrogen peroxide.(11) This suggested that HSPB1 could be protecting IEC during ethanol stress by supporting activation of the Nrf2 transcriptional program. Interestingly, Nrf2 transcriptional activity was not only activated in HSPB1 deficient cells during ethanol exposure, it was also much higher (approximately 3-fold) in the KO cells than WT cells (**Fig. 4A**). To verify that compromised Nrf2 activity was not responsible for the increased death we observed in the KO cells, we immunoblotted for two key Nrf2 target genes, GPX4 (glutathione peroxidase 4) and FSP1 (ferroptosis suppressor protein 1) in ethanol stressed cells. GPX4 and FSP1 act in parallel to protect cells from ferroptosis. Levels of both proteins were similar in WT and KO cells, both before and after ethanol challenge (**Fig. 4B**).

**Figure 4.**
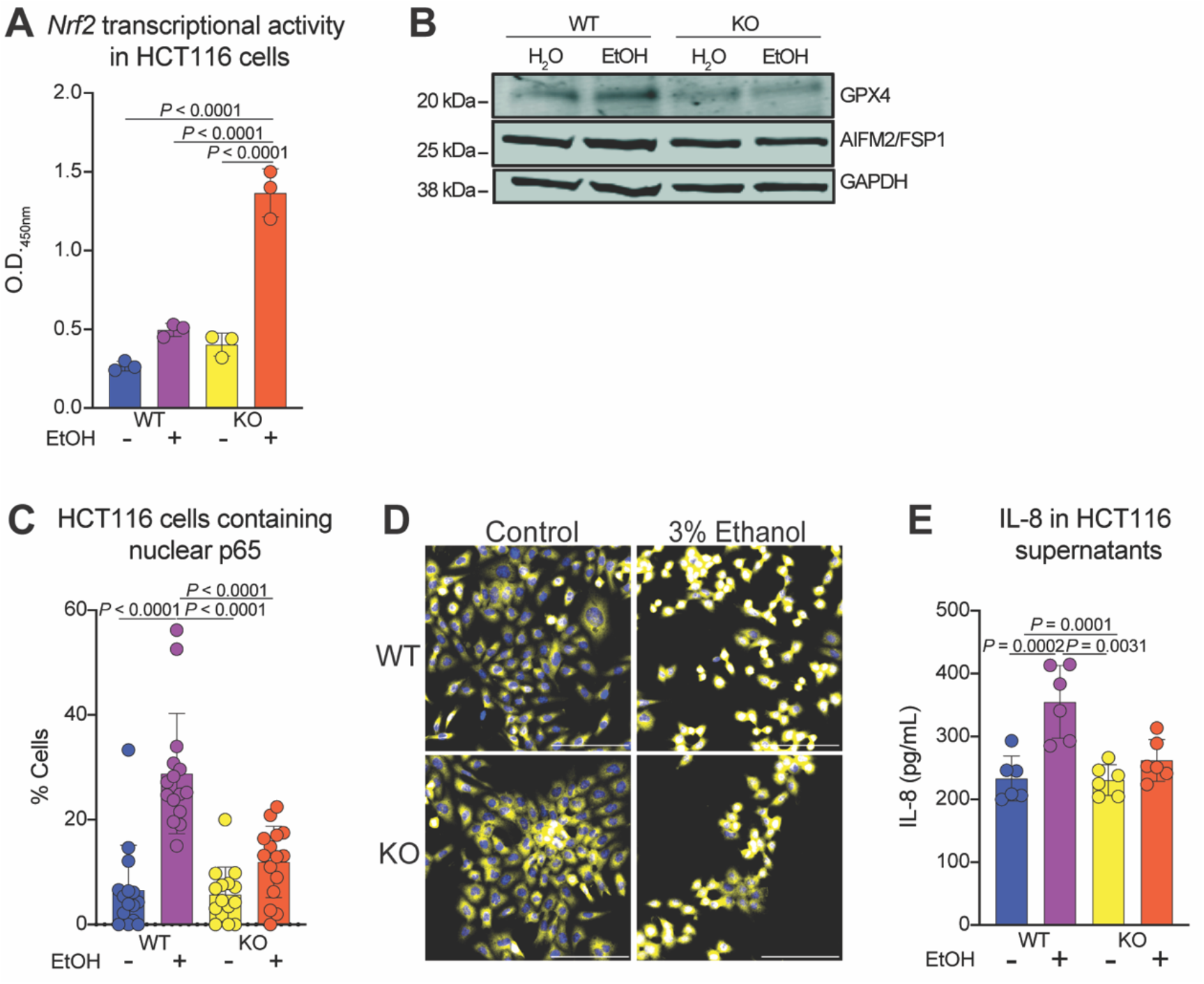
HSBP1 is not required for activation of Nrf2 transcriptional activity, but is necessary for NF-_κ_B activation during ethanol stress. **A.** Nrf2 transcriptional activity measured in WT or KO HCT116 cells treated with media or 1.5% ethanol for 1 hour. **B**. Immunoblot for GPX4, AIFM/FSP1 and GAPDH in WT and KO HCT116 cells exposed to media or 1.5% ethanol for 1 hour. **C**. Percentage of WT or KO HCT116 cells containing nuclear NF-kB p65 after treatment with media or 1.5% ethanol for 30 minutes. **D**. Immunofluorescence of WT or KO HCT116 cells stained with NF-kB p65 (yellow) and Hoechst (blue) after treatment with media or 1.5% ethanol for 30 minutes. Scale bar = 100 µm, 400X magnification. **E**. IL-8 production in WT or KO HCT116 cells treated with media or 1.5% ethanol overnight. Data plotted as mean + SD. (A,C,E) One-way ANOVA with a Tukey multiple comparisons post hoc test.

NF-kB (Nuclear factor kappa-light-chain-enhancer of activated B cells) is a second transcription factor that coordinates cellular defense against oxidative damage.(31) Epithelial NF-kB signaling directly suppresses programmed death pathways through activation of pro-survival genes. Additionally, complex cross-talk between NF-kB and Nrf2 maintains balanced inflammation to prevent activation of programmed cell death pathways.(32) Intracellular HSPB1 activates NF-kB through direct interaction with IκBα (the inhibitory protein of NF-κB) and promotion of its degradation, which allows NF-kB to translocate to the nucleus.(33) Extracellular HSPB1 activates NF-kB by binding to and triggering pattern recognition receptors (PRR), including TLR2 (toll-like receptor 2), TLR4, TLR5, and TLR7.(34) Translocation of the NF-kB component p65 to the nucleus was similar in WT and KO cells at baseline, but the percentage of cells with nuclear p65 was lower in KO cells after exposure to ethanol (**Fig. 4C, D**). Likewise, levels of the NF-kB regulated chemokine CXCL8 (C-X-C motif chemokine ligand 8), commonly known as Interleukin-8 (IL-8), were lower in KO cells than WT cells exposed to ethanol (**Fig. 4E**). Together, these data show that HSPB1 supports NF-kB activation, but is dispensable for Nrf2 activity, in IEC stressed by ethanol.

### Extracellular HSPB1 protects IEC from ethanol activated death

Intracellular HSPB1is thought to limit ferroptosis through regulation of signaling cascades that reduce iron accumulation in stressed cells, stabilization of the cytoskeleton to decrease iron import into cells, and inhibition of ferritin degradation to sequester free iron.(35) No roles for extracellular HSPB1 have been described in ferroptosis. However, our previous study demonstrating that hepatocytes release HSPB1 in response to ethanol combined with our data that HSPB1 participates in NF-kB activation during ethanol stress led us to consider an extracellular role for HSPB1 in cell survival during ethanol exposure. We first determined whether IEC, like hepatocytes, release HSBP1 in response to ethanol. Similar to what we found with intracellular HSPB1, extracellular HSPB1 increased with ethanol exposure and continued to increase with repeated exposure over several days (**Fig. 5A**). Next, we considered whether extracellular HSPB1 could protect IEC from ethanol-activated programmed cell death mechanisms. We found that treatment with recombinant HSPB1 (rHSPB1) prior to ethanol challenge protected KO cells from death (**Fig. 5B**). To ensure that the protective effect of extracellular HSPB1 in ethanol stress was not cell type specific, we measured HSPB1 released from a second IEC line during ethanol stress. As with HCT116 cells, Caco2 cells released HSPB1 in response to ethanol (**Fig. 5C**). Likewise, administration of rHSPB1 protected Caco2 cells from ROS generated by ethanol exposure, even though these cells are able to release HSPB1 themselves (**Fig. 5D**). Presumably, this is due to the fact that with rHSPB1 administration, HSPB1 does not have to be produced and released by these cells a priori, it is already present in the extracellular environment and activating the NF-kB-mediated antioxidant response. Thus, extracellular HSPB1 is responsible for limiting ROS and cell death in IEC exposed to ethanol.

**Figure 5.**
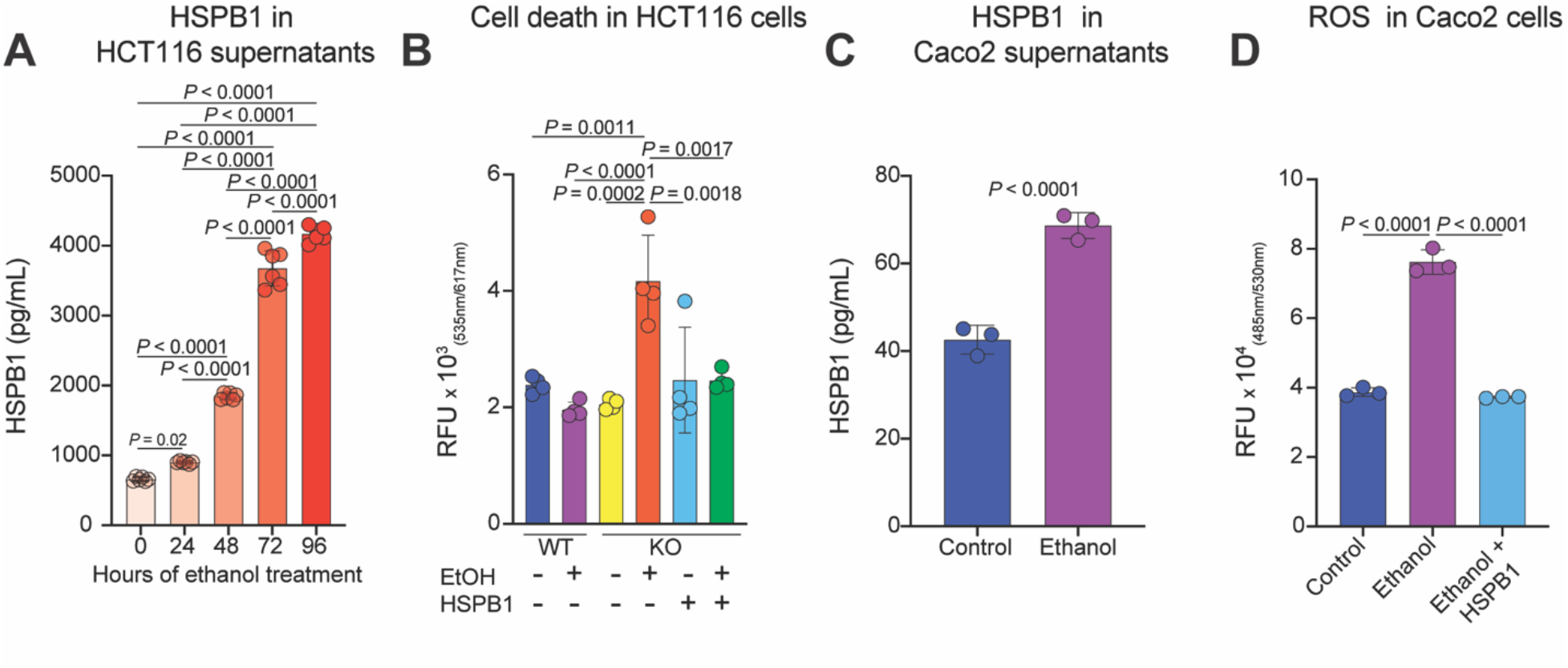
Extracellular HSPB1 protects IEC from ethanol activated death. **A.** HSPB1 in media of WT HCT116 cell stimulated with 1.5% ethanol every 24 hours for 96 hours. **B**. Cell death in WT or KO HCT116 cells treated with rHSPB1for 1 hour prior to stimulation with media or 3% ethanol for 2 hours measured by PI. **C**. HSPB1 in media from Caco2 cells stimulated with media or 1.5% ethanol for 1 hour. **D**. Reactive oxygen species measured in Caco2 cells treated with 3 µM rHSPB1 prior to media or 1.5% ethanol for 1 hour. Data plotted as mean + SD. (A,B,D) One-way ANOVA with a Tukey multiple comparisons post hoc test. (C) Unpaired *t* test.

## DISCUSSION

Acute ethanol toxicity is a consequence of alcoholic beverage consumption that affects organs throughout the body, perhaps none more than the GI tract since it is the first point of contact with high concentrations of ingested alcohol. In this study, we report that HSPB1 is upregulated in and released from IEC experiencing ethanol stress. HSPB1 supports NF-kB activity in ethanol exposed IEC and limits their death by ferroptosis. Recombinant HSPB1 is sufficient to rescue IEC from ethanol activated death mechanisms, indicating that extracellular HSPB1 is responsible for its protective function in this context. These findings support the previously identified role for HSPB1 as a damage-associated molecular pattern (DAMP) in ethanol toxicity. DAMPs are released from stressed or damaged cells and activate inflammation in surrounding cells. The majority of research into DAMPs has focused on their detrimental effects. However, inflammation can also promote survival in stressed cells, as we found in this study. This suggests a more nuanced role for HSPB1 in ethanol activated inflammation than previously understood.

Ethanol activated inflammation has primarily been studied as a self-damaging disease mechanism in chronic alcohol consumption. However, inflammation also plays important roles in tissue homeostasis in the gut. For example, MAMPs released from gut microbes maintain a low level of NF-kB activity in IEC and inhibition of NF-kB signaling causes spontaneous epithelial cell death and colitis in mice.(36-38) This has led to the understanding that NF-kB dependent inflammation is delicately balanced in the gut. It is thought that low levels are required to maintain epithelial barrier health, while high or sustained levels of NF-kB activity lead to self-damaging inflammation and immune activation. In addition to its well-characterized inflammatory functions, the NF-kB signaling pathway regulates ferroptosis through transcriptional activation of at least two distinct mechanisms.(39, 40) In the first, NF-kB activates antioxidant pathways that increase cysteine uptake and glutathione biosynthesis to counteract oxidative stress. In the second, NF-kB activates genes that encode iron-scavenging or sequestration mechanisms to limit concentrations of free iron in cells. Previous studies have demonstrated that intracellular HSPB1 regulates NF-kB activity through direct interaction with IκBα and that extracellular HSPB1 regulates NF-kB activity through activation of toll-like receptors on the cell surface. Our finding that extracellular HSPB1 is sufficient to protect IEC from ethanol activated death suggests that ethanol stressed cells release HSPB1 to signal danger to neighboring cells and cause them to proactively upregulate antioxidant systems in anticipation of oxidative damage. It also suggests that HSPB1 is acting as a DAMP and signaling through PRR to execute this protective action. This may be particularly important in the gut since some resident gut microbes can produce ethanol, leading to physiologic exposure to this toxicant.(2, 41) Ultimately, extracellular HSPB1 is most likely to play an important role in protecting the gut epithelium during physiologic ethanol exposure or acute ethanol exposure through alcoholic beverage consumption.

The level of HSPB1 protein in the intestinal epithelium is calibrated in response to inflammatory cytokines delivered from innate and adaptive immune cells underlying the intestinal epithelium and microbial signals from the gut lumen.(9, 10, 14, 15, 42) In the normal small intestine, cytokine and microbial signals are low, leading to low to non-detectable levels of HSPB1 protein within IEC. In this study we found that cultured IEC increase intracellular HSPB1 levels rapidly and release HSPB1 in response to ethanol. We primarily utilized colorectal cancer cell lines for this study. Transformed IEC constitutively express HSPB1, which is thought to perform an intracellular, cytoprotective role in these cells.(43) It is possible that this high level of constitutive HSPB1 expression caused this model to overestimate the amount of HSPB1 released from IEC in response to ethanol. Although our findings in transformed IEC were similar to our findings in intestinal tissue challenged with ethanol *ex vivo*. It is also possible that the constitutive HSPB1 expression caused the impact of extracellular HSPB1 protection to be underestimated by this model. For these reasons, it will be important to validate our findings in follow-up studies utilizing primary cells and more complex intestinal modeling.

Together, our findings directly link HSPB1 to intestinal cytoprotection in acute ethanol toxicity and demonstrate that extracellular HSPB1 is responsible for this function. This was unexpected since an extracellular HSPB1function has not been demonstrated in the gut. However, extracellular HSPB1 has been shown to be protective during ethanol stress in other cell types. We propose that rapid upregulation and release of HSPB1 is a widespread response to ethanol stress that is protective in acute ethanol exposure. However, when extracellular HSPB1 levels are sustained due to chronic alcohol consumption, they contribute to self-damaging inflammation. The data generated in this study and previous literature support this model, but key questions remain unanswered. For example, does HSPB1 released from IEC stay local or translocate out of the gut? We previously demonstrated that HSPB1 is increased in the blood and liver of patients with alcohol-associated hepatitis, a highly inflammatory and often life-threatening form of alcohol-associated liver disease.(16) HSPB1 protein levels were also increased in the liver in a murine model of chronic alcohol consumption and hepatocytes released HSPB1 in response to ethanol. This extracellular HSPB1 triggered TNFα release from macrophages, suggesting that extracellular HSPB1 contributes to self-damaging inflammation in alcohol-associated hepatitis. This means that the release of HSPB1 from the small intestinal epithelium could be particularly problematic in chronic alcohol consumption if it translocates to the liver through the portal circulation and contributes to inflammation there. It is also unclear which triggers of HSPB1 production and release are most important in the gut during acute ethanol toxicity. Ethanol itself can trigger HSPB1 release from IEC, but it can also damage gut microbes, leading to release of MAMPs that activate HSPB1 expression in IEC. In summary, our data from this and previous studies suggest that HSPB1 is essentially a “double edged sword” in the context of ethanol stress. By this we mean that HSPB1 plays critical cytoprotective roles during acute ethanol stress, but can lead to self-damaging inflammation in patients chronically consuming alcohol. Therefore, novel diagnostic and therapeutic strategies designed to limit dangerous levels of extracellular HSPB1 along the gut-liver axis have the potential to identify patients at risk of severe inflammatory disease and short-circuit inflammation to protect ethanol-stressed organs from life-threatening injury.

## METHODS

### Ex vivo ethanol treatment of the small intestine

All animal experiments were approved by the Cleveland Clinic IACUC (Protocol #2595). Specific pathogen free C57BL/6 mice were euthanized and the proximal 10 cm of small intestine was collected immediately post-mortem. The small intestine was tied off in two 5 cm segments using sterile suture. One segment was filled with vehicle (sterile water) using a gavage needle attached to a 1 ml syringe and then the open end of the intestine was tied off with sterile suture. The other segment received 5% ethanol in the same manner. The tissue was incubated in Dulbecco’s Modified Eagle Medium (DMEM) with 10% fetal bovine serum (FBS) for 30 min. at 37°C and 5% CO_2_. After incubation, the tied ends were cut, the sample was placed in 500 µL of cell lysis buffer (Cell Signaling, 9803S) supplemented with protease inhibitor (Roche, 11836170001), and the tissue was homogenized on ice using a Tissue-Tearor (BioSpec). After a 15-minute incubation, samples were centrifuged at 10,000 xg for 10 minutes at 4°C and the supernatants were collected for ELISA.

### HSPB1 ELISA

Samples (volume normalized by protein concentration) were loaded in duplicate or triplicate onto a high-binding polystyrene plate (Costar, 07-200-39) and incubated overnight at 4°C. Recombinant human HSPB1 was used as the protein standard. The following day, lysate was decanted, and the plate washed once with 200 µL per well of PBS with 0.01% Tween-20. The plate was blocked using 1% bovine serum albumin (BSA) in phosphate buffered saline (PBS; 200 µL per well) for 1.5 hours at room temperature. After blocking, 50 µL of 0.67 µg/mL anti-HSPB1 antibody (custom antibody produced by Biomatik, AB006678) was added to the plate and incubated for 2 hours at room temperature. After primary antibody incubation, the plate was washed 3 times with 200 µL of PBS with 0.01% Tween-20 per well. Then 50 µL of secondary antibody conjugated with horseradish peroxidase (Jackson Immunoresearch, 711-035-152) was added to the plate at a final concentration of 0.08 µg/mL for 1 hour. The plate was then again washed 3 times using PBS with 0.01% Tween-20 and 50 µL of pre-warmed TMB (3,3’,5,5’-tetramethylbenzidine) was added to the plate. The detection reagent was developed for 5 minutes at room temperature and then the reaction was quenched with 50 µL of 0.5 M sulfuric acid. Plates were read using a BioTek Synergy HTX multimode reader.

### Immunoblotting

Protein concentration was quantified using the Pierce BCA assay kit (Thermo Scientifc). 30 μg of sample was subjected to SDS-PAGE under reducing conditions on a 4-12% Bis-Tris gel (Invitrogen) with NuPAGE MOPS buffer (Invitrogen). Protein was transferred to Immobilon-FL PVDF membrane with 0.45μm pore size (Millipore) and blocked with Intercept (PBS) blocking buffer (LICORbio). After blocking, the membrane was cut at 30kDa and the two membranes were incubated with either 0.67 μg/mL rabbit anti-HSPB1 polyclonal antibody (Biomatik, AB006678) or 0.5 μg/mL rabbit anti-AIM2/ FSP1 polyclonal antibody (Proteintech, 20886-1-AP) overnight at 4°C with agitation. The following day, membranes were washed 3 times in PBS with 0.01% Tween-20 for 5 minutes each wash and then incubated with 0.05 μg/mL IRDye800CW donkey anti-rabbit secondary antibody (LICORbio, 926-32213) in blocking buffer for one hour. Membranes were then again washed 3 times in PBS with 0.01% Tween-20 for 5 minutes and fluorescent signal was detected using an LI-COR Odyssey CLX machine and LICOR Acquisition software. Membranes were then stained with 1 μg/mL mouse anti-GPX4 monoclonal [3F5G5] antibody (ProteinTech, 67763-1-Ig) overnight at 4°C with agitation. The following day, membranes were washed 3 times in PBS with 0.01% Tween-20 for 5 minutes each wash and incubated with 0.05 μg/mL IRDye680RD donkey anti-mouse secondary antibody (LICORbio, 926-32212) in blocking buffer for one hour. Membranes were then again washed 3 times with PBS with 0.01% Tween-20 for 5 minutes and fluorescent signal was detected using an LI-COR Odyssey CLX machine and LICOR Acquisition software. Densitometry was performed using ImageJ.

### NF-kB p65 localization in HCT116 cells

WT and KO HCT116 cells were plated on coverslips at 0.1 × 10^6^ cells/well in a 12-well tissue culture plate. The following day cells were washed once with PBS and 1.5% ethanol in DMEM or DMEM only was added for 30 minutes with cells incubated at 37°C with 5% CO2. Following incubation cells were washed once with PBS, 4% paraformaldehyde (PFA) was added, and cells were incubated at room temperature for 10 minutes. Cells were washed three times with PBS and permeabilized with 0.1% TritonX-100 in PBS for 10 minutes at room temperature. Cells were washed three times with PBS then blocked with Superior blocking buffer (G-Biosciences, C752H38) for 60 minutes at room temperature. Cells were stained with 0.35 µg/mL of NF-kB p65 (Cell Signaling, 82425) overnight. The following day the cells were washed three times and stained with 1 µg/mL Alexa Fluor 647 donkey anti-rabbit (Jackson Immunoresearch, 711-605-152) secondary antibody for 2 hours. Coverslips were counterstained using 10 μg/mL bisBenzimide H 33258 dissolved in TBS for 20 minutes in the dark at room temperature then mounted on slides using ProLong glass antifade mounting media (Thermo Fisher, P36980). Coverslips were imaged at a 400X magnification using a BioTek Lionheart FX automated microscope. All images were viewed and edited in the Gen5 3.18 software. Gen5 3.18 software was used to quantify the total number of cells per image based on the signal in the DAPI channel (Hoechst). A mask was created to measure and filter the intensity of the Cy5 (p65) signal and identify the subpopulation of cells with nuclear p65 based on the intensity of the signal within the nuclear region of interest identified. To determine the percentage of cells with nuclear p65, the number of cells in the subpopulation with intense nuclear p65 signal was divided by the total number of cells.

### Cell Based Assays

#### HSPB1 measured in HCT116

(1B and 4B) WT and KO HCT116 cells were plated at 2.5 × 10^4^ cells/well in a 6-well tissue culture plate. The following day, cells were treated with 1.5% ethanol in serum-free McCoy’s 5A media for 1 hour at 37°C with 5% CO2. Cells were lysed in 125 µL RIPA lysis buffer supplemented with protease inhibitor (Roche, 11836170001). After a 15-minute incubation, samples were centrifuged at 12,000 xg for 10 minutes at 4°C and the supernatants were collected for immunoblot. (1D) WT HCT116 cells were plated at 2.5 × 104 cells/well in a 6-well tissue culture plate. The following day, the cells were treated with fresh 1.5% ethanol in serum-free McCoy’s 5A media every 24 hours for 72 hours incubated at 37°C with 5% CO2. Cells were lysed in 300 µL cell lysis buffer (Cell signaling, 9803S) supplemented with protease inhibitor (Roche, 11836170001). After a 15-minute incubation, samples were centrifuged at 12,000 xg for 10 minutes at 4°C and the supernatants were collected for ELISA. Media only controls were included in all experiments.

#### Cell Death measured using propidium iodide

WT and KO HCT116 cells were seeded at 1.0 × 10^4^ cells per well in a black-walled, clear bottom 96-well tissue culture plate. The following day the cells were washed once with PBS and treated with either phenol red free (PRF) DMEM or 3% ethanol in PRF DMEM. After a two-hour incubation at 37°C with 5% CO2, 30 ug/mL of propidium iodide (PI) was added to each well and incubated for 30 minutes at 37°C with 5% CO2. Plates were read using a BioTek Synergy HTX multimode reader. Plates were then imaged at a 40X magnification on the BioTek Lionheart FX Automated Microscope. All images were captured using the TRITC (PI) channel, and images were viewed and edited in the Gen5 3.18 software.

#### HCT116 cells incubated with 1.5% ethanol followed by 3% ethanol shock

WT and KO HCT116 cells were plated at 0.1 × 10^6^ cells/dish in a 35mm glass bottom dish. The following day, cells were exposed to 1.5% ethanol overnight and then 3% ethanol for 1 hr. Cells were incubated at 37°C and 5% CO_2_ for the duration of the experiment. After ethanol treatment, cells were rinsed once with PBS and the LIVE/DEAD® Viability/Cytotoxicity Kit for mammalian cells (Invitrogen, # L3224) was used according to the manufacturers protocol. In short, 150 µL of the LIVE/DEAD reagents were added to the glass bottom dish and incubated at room temperature for 30 minutes. The excess dye was removed and replaced with 150 µL of PBS, the cells were scraped into a 96 well plate and read using a BioTek Synergy HTX multimode reader.

#### ROS (DCFDA) measurement in ethanol treated HCT116

WT and KO HCT116 cells were plated at 0.1 × 10^6^ cells/dish in a 35mm glass bottom dish. The following day, cells were exposed to 1.5% ethanol for 1 hour at 37°C and 5% CO_2_. After treatment, cells were rinsed once with PBS. 2′,7′-Dichlorodihydrofluorescein diacetate (DCFDA) in serum-free McCoy5a media was diluted to a working concentration of 10 µM and added to the glass bottom dish. Cells were incubated for 30 minutes 37°C with 5% CO2. Cells were scraped into a 96 well plate and read using a BioTek Synergy HTX multimode reader.

#### Measurement of Protein Carbonylation in ethanol treated HCT116

WT and KO HCT116 cells were plated at 2.5 × 10^4^ cells/well in a 6-well tissue culture plate. The following day, the cells were treated with 1.5% ethanol for 1 hour at 37°C and 5% CO_2_. After ethanol treatment, cells were rinsed once with PBS. Cells were then scraped from each well into 50 mM MES 1 mM EDTA buffer and sonicated for 1 minute on ice to ensure complete lysis. Lysate was centrifuged for 15 minutes at 10,000 X G at 4°C and the protein carbonylation assay kit (Cayman Chemical Company, 10005020) was used per the manufacturer’s instructions.

#### Measurement of Iron Accumulation (FerroOrange) in ethanol treated HCT116

WT and KO HCT116 cells were plated at 5.0 × 10^4^ cells/well in a clear 96-well tissue culture plate. The following day, cells were treated with 1.5% ethanol for 1 hour at 37°C and 5% CO_2_. After ethanol treatment the cells were rinsed once with PBS and 1 μM/L FerroOrange (Dojindo, # F374) working solution was added to the cells and incubated for 30 min at 37°C and 5% CO2. After incubation, the plate was read using a BioTek Synergy HTX multimode reader.

#### Nrf2 transcriptional activity in ethanol treated HCT116 cells

WT and KO HCT116 cells were plated at 2.5 × 10^4^ cells/well in a 6-well tissue culture plate. The following day, the cells were treated with 1.5% ethanol for 1 hour at 37°C and 5% CO_2_. Lysates were collected and analyzed using the Nrf2 transcriptional factor assay kit (Cayman Chemical, 600590) according to the manufacturer’s instructions.

#### IL-8 production in ethanol treated HCT116 cells

WT and KO HCT116 cells were plated at 0.3 × 10^6^ cells/well in a 6-well tissue culture plate. The following day, cells were rinsed once with PBS and treated with 1.5% ethanol in serum-free McCoy’s 5A media and incubated overnight at 37°C and 5% CO_2_. Supernatants were collected and analyzed using the Human IL-8/CXCL8 DuoSet ELISA (R&D systems, DY208) according to the manufacturer’s instructions.

#### HSPB1 production in ethanol treated HCT116 cells cycled with 1.5% ethanol for 96 hours

WT HCT116 cells were plated at 0.3 × 10^6^ cells/well in a 6-well tissue culture plate. The following day, cells were rinsed once with PBS and treated with 1.5% ethanol in serum-free McCoy’s 5A media and incubated overnight at 37°C and 5% CO_2_. Media was replaced with fresh 1.5 % ethanol every 24 hours for 96 hours. Supernatants were collected for ELISA.

#### Cell death in cells treated with rHSPB1 measured using propidium iodide

WT and KO HCT116 cells were seeded at 1.0 × 10^4^ cells per well in a clear 96-well tissue culture plate. The following day the cells were washed once with PBS and treated with either DMEM alone or 3 µM rHSPB1 for 1 hour and then 3% ethanol was added for 3 hours. Recombinant HSPB1 was produced as previously described.(16) Cells were incubated at 37°C and 5% CO2 for the duration of the experiment. Following incubation, 30 ug/mL of PI was added to each well and incubated for 30 minutes at 37°C and 5% CO2. Plates were read using a BioTek Synergy HTX multimode reader.

#### HSPB1 production in ethanol treated Caco2 cells treated with 1.5% ethanol

WT Caco2 cells were plated at 0.3 × 10^6^ cells/well in a 6-well tissue culture plate. The following day, cells were rinsed once with PBS and treated with 1.5% ethanol in serum-free DMEM media for 1 hour at 37°C and 5% CO_2_. Supernatants were collected for ELISA.

#### ROS (DCFDA) measurement in ethanol treated Caco2 cells

WT Caco2 cells were plated at 0.1 × 106 cells/dish in a 35mm glass bottom dish. The following day, cells were treated with 1.5% ethanol for 1 hour and 3 µM HSPB1 1 hr and incubated at 37°C and 5% CO_2_. After treatment, cells were rinsed once with PBS. 2′,7′-Dichlorodihydrofluorescein diacetate (DCFDA) in serum-free McCoy5a media was diluted to a working concentration of 10 µM and added to the glass bottom dish. Cells were incubated for 30 minutes 37°C with 5% CO2. Cells were scraped into a 96 well plate and read using a BioTek Synergy HTX multimode reader.

#### Statistical analysis

All cellular experiments were replicated independently, at least 3 times. Values shown in all figures represent the means ± SD. Pairwise differences were evaluated using parametric 2-tailed *t-*tests. Differences among multiple groups were detected using 1-way ANOVA with Tukey post hoc analysis. Statistical analysis was performed using GraphPad Prism (v 10.6). A *p* value of <0.05 was considered significant.

## Author Contributions

Conceptualization: JSM. Methodology: JSM, AMCO, MB. Investigation: MB, AMCO, BBK, SO, KAH. Funding acquisition: JSM. Project administration: JSM. Supervision: JSM, AMCO. Writing – original draft: JSM. Writing – review & editing: JSM, AMCO, MB, BBK, SO, KAH

## Data and materials availability

All data used in the analysis are available upon request. Materials used in this study are available upon request and with completion of the appropriate material transfer agreements.

